# Towards a study of gene regulatory constraints to morphological evolution of the *Drosophila* ocellar region

**DOI:** 10.1101/031948

**Authors:** D. Aguilar-Hidalgo, D. Becerra-Alonso, D. García-Morales, F. Casares

## Abstract

The morphology and function of organs depend on coordinated changes in gene expression during development. These changes are controlled by transcription factors, signaling pathways and their regulatory interactions, which are represented by *gene regulatory networks* (GRNs). Therefore, the structure of an organ GRN restricts the morphological and functional *variations* that the organ can experience –its potential morphospace. Therefore, two important questions arise when studying any GRN: what is the predicted available morphospace and what are the regulatory linkages that contribute the most to control morphological variation within this space. Here, we explore these questions by analyzing a small “3-node” GRN model that captures the Hh-driven regulatory interactions controlling a simple visual structure: the ocellar region of *Drosophila.* Analysis of the model predicts that random variation of model parameters results in a specific non-random distribution of morphological variants. Study of a limited sample of Drosophilids and other dipterans finds a correspondence between the predicted phenotypic range and that found in nature. As an alternative to simulations, we apply Bayesian Networks methods in order to identify the set of parameters with the largest contribution to morphological variation. Our results predict the potential morphological space of the ocellar complex, and identify likely candidate processes to be responsible for ocellar morphological evolution using Bayesian networks. We further discuss the assumptions that the approach we have taken entails and their validity.

## INTRODUCTION

The evolution of animals has resulted in a staggering diversity of forms. But what are the limits to morphological variation? The answer to this question requires considering that the shape of body parts is controlled by complex genetic programs operating during embryonic development. These programs integrate the action of many genes across growing fields of cells forming extensive developmental *gene regulatory networks* (“GRN”) (Arnone and Davidson, 1997). Therefore, if form is determined to a large extent by gene networks, it follows that these networks should restrict the potential evolutionary routes to morphological variation (Oster et al., 1988; Kauffman, 1993; Arthur, 2006; Davidson and Erwin, 2006; Felix, 2012; Jaeger and Monk, 2014), an idea first formulated by C. H. Waddington (Waddington, 1957). Determining the potential range of phenotypes allowed by a particular GRN, however, is not straightforward, because gene networks are complex and their analysis often entails the combined use of model organisms and mathematical simulations. Examples of this combined approach in animal development are studies analyzing the contribution of gene network organization (or topology) to morphological variation of teeth (Salazar-Ciudad and Jernvall, 2010; Harjunmaa et al., 2014), the number and pattern of digits in the tetrapod limb (Lopez-Rios et al., 2014; Raspopovic et al., 2014), the patterning of the *Drosophila* eggshell epithelium (Faure et al., 2014) or the segmentation of the early *Drosophila* embryo ((Jaeger et al., 2004); see also (Felix, 2012) for a recent review).

Another experimental system well suited to study the relation between a developmental GRN and morphological variation is the ocellar region in dipterans. The ocellar region is part of the visual system of insects and is morphologically simple: it comprises three single-lens eyes (the ocelli) located at the vertices of a triangular cuticle patch on the insect dorsal head (Figure 1A). Therefore, main quantitative traits in this system are the sizes of the ocelli and their separating (“interocellar”) distance. Interestingly, the ocellar region shows morphological variation in different fly species (Figure 1C,D), which permits to explore not only the phenotypic variation induced experimentally in one model organism *(D. melanogaster)*, but also the variation generated during evolution across species. Recently, our group generated a GRN model of the ocellar region patterning (Aguilar-Hidalgo et al., 2013). In this GRN, the evolutionary conserved Hedgehog (Hh) signaling pathway plays a pivotal role, controlling the specification of the two major fates (retina/ocellus and interocellar cuticle), as well as their size and spacing (Royet and Finkelstein, 1996; Royet and Finkelstein, 1997; Blanco et al., 2009; Brockmann et al., 2011; Aguilar-Hidalgo et al., 2013; Dominguez-Cejudo and Casares, 2015) (Figure 1B). One of the most interesting predictions derived from this GRN model was that random variations in parameter sets resulted in a non-random specific morphological space. Therefore, one potential application of the GRN model analysis could be the identification of the parameters controlling the paths to morphological variation within this restricted space. However, the GRN model in Aguilar-Hidalgo (2013) was very complex (1 partial differential equation and 12 ordinary differential equations with 68 parameters, of which 32 were studied) which makes this sort of analysis cumbersome.

**Figure 1.**
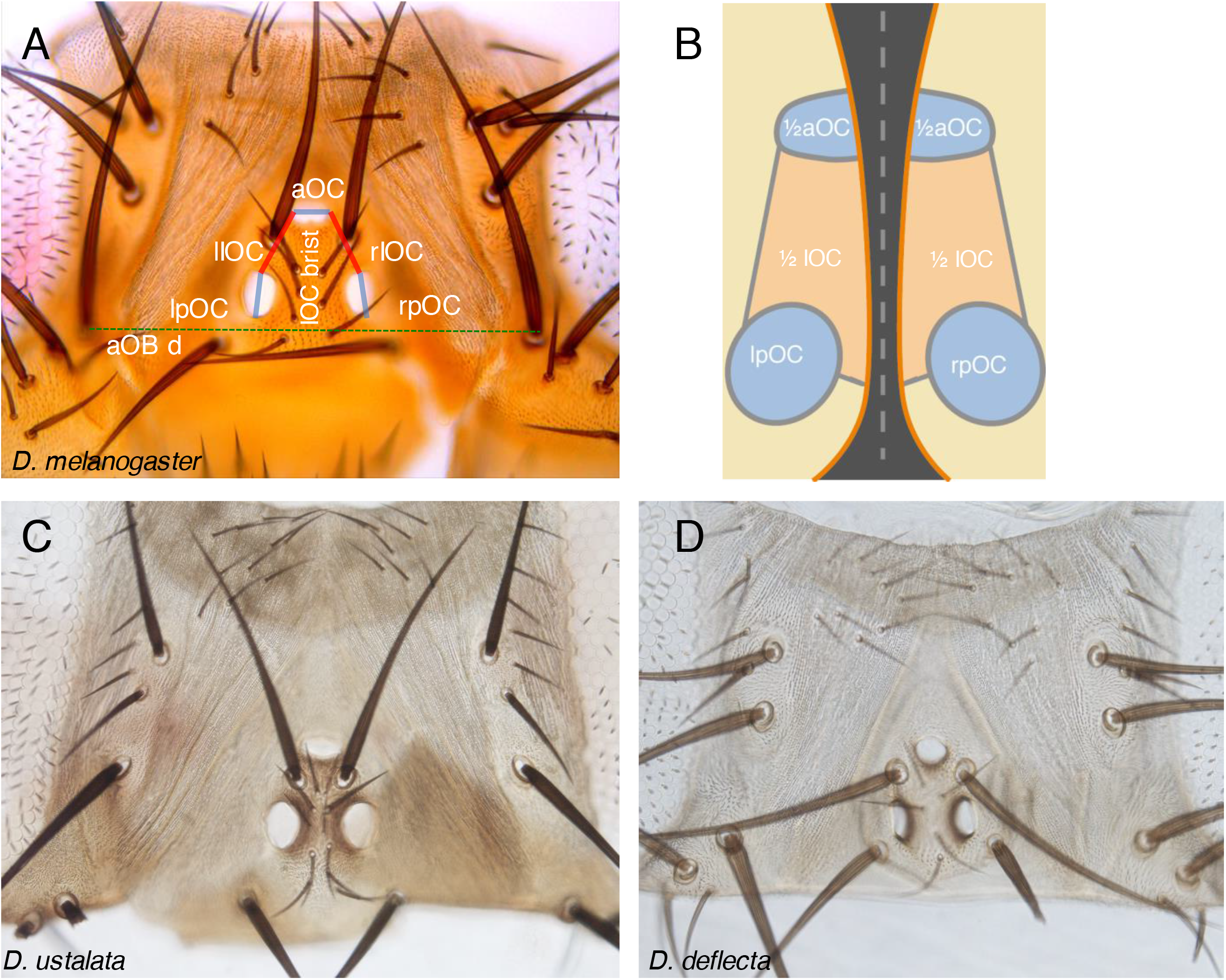
Ocellar region structure. (A) Dorsal view of a *Drosophila melanogaster* female head. Anterior ocellus (“aOC”) and posterior ocelli (left and right “pOC”). Left and right interocellar distances (“rIOC” and “lIOC”) are marked in red. OC lengths are marked in blue. The distance between the anterior orbital bristles (“aOBd”, dashed green line) is used to normalize for head size. “IOC brisl.”: interocellar bristles. (B). Schematic representation of the dorsal fusion of the left and right head primordia. The aOC and the interocellar regions are formed by the fusion of the contralateral halves. The pOC remain separate. (C, D) Dorsal heads from *D. ustalata* and *D. deflecta* female adults, to scale. OC and IOC length differ.

Here, we used a reduced “three-node” GRN that still recapitulates the expression patterns of key genes in the ocellar region. Our results indicate that the topology of the ocellar GRN defines a particular potential morphological space for the ocellar complex. In this GRN, quantitative changes in parameter values seem sufficient to explain the quantitative morphological variation found in nature without the need of gene network rewiring. Our analysis further identifies likely candidate processes to be responsible for ocellar morphological evolution.

## MATERIALS AND METHODS

### Fly species and *D. melanogaster* strains

*Drosophila melanogaster* (strain Oregon-R). *D. gunungcola, D. lutescens, D. lulchrella, D. guttifera, D. prolongata, D. ustulata, D. deflecta, D. fuyamai, D. suzukii, D. biarmipes, D. pseudoobscura, D. bipectinata, D. ananassae, D. sechellia, D. mauritiana, D. yakuba, D. parabipectinata, D. kikkawai, D. teissieri, D. santomea, D. takahashii, D. eugracilis, D. simulans, D. orena, D. erecta, D. willistoni, Chymomyza pararufithorax* were obtained as EtOH-preserved specimens from B. Prud’homme (IBDML, Marseille); *D. virilis* from J. Vieira (IBMC/I3S, Oporto); *Megaselia abdita* and *Episyrphus balteatus* (EtOH-preserved) from J. Jeager/K. Wotton (CRG, Barcelona/KLI, Vienna); *Calliphora vicina* from P. Simpson (U. Cambridge, Cambridge); *Ceratitis capitata* and *Bactrocerus oleae* (EtOH-preserved) from M. Averof (IGFL, Lyon). *D. hydei* (strain KS13) was established as a culture at the CABD (Seville). *Megaselia scalaris* specimens were captured at the CABD fish facility; *Musca domestica* and other dipteran specimens were captured from the wild. The phylogenetic range of this collection spans about 150 Million years (Myrs), with Phoridae *(M.abdita* and *M.scalaris)* having the oldest origin. The divergence time of Syrphidae has been set about 95 Myrs ago. The remaining species belong to Schizophora, with an estimated origin 75 Myrs ago (for an updated and detailed Dipteran phylogeny, please check (Wiegmann et al., 2011).

In addition, the following *D. melanogaster* strains were used: *en-Z (en[xho25];* Flybase: FBti0002246); a *hh-GAL4, UAS-GFP::Hh* strain was used to monitor the Hh expression domain in the ocellar complex (Callejo et al., 2008).

### Head cuticle preparation and measurements

Dorsal head cuticle pieces were dissected from adult or late female pharate heads in PBS, and mounted in Hoyers’ solution:acetic acid (1:1), as described in (Casares and Mann, 2000). Images were obtained in a Leica DM500B microscope with a Leica DFC490 digital camera. Measurements were carried out using the line measurement tool of ImageJ (Rasband, 1997–2014).

### Immunostaining and imaging

Immunofluorescence in eye imaginal discs and embryos was carried out according to standard protocols. Antibodies used were: mouse anti-Eya (10H6; from Developmental Studies Hybridoma Bank, University of Iowa (http://dshb.biology.uiowa.edu/) 1/200; rabbit anti-ß-galactosidase antibody (Cappel), 1/1000; mouse anti-Ptc (gift from I. Guerrero, CBM-SO, Madrid), 1/100; rabbit anti-GFP (A11122, Molecular Probes), 1/1000. Alexa-conjugated anti-Rabbit-488 and anti-mouse-555 secondary antibodies were used at a 1/1000 dilution. Image acquisition was carried out in a Leica SP2 AOBS confocal microscope. Images were processed with Adobe Photoshop CS5.

### Model simulation

To simulate the 3node-GRN ocellar region model we first assume that the Hh profile is in steady state. We can assume this as we want to compare signaling patterns with sizes of differentiated tissues in adult flies, thus the development of the ocellar region is in steady state. Additionally, we do not consider tissue growth, but instead the Hh profile grows in a fixed-size grid. We solved the reaction-diffusion equation for Hh (equation S1) in steady state analytically, the solution of which is a spatial dependent function Hh(x) (equation 1). This function serves as input to the three ordinary differential equations that show the spatial pattern for PtcHh, CiA and En (see equations 2-4 in Fig. 2D). Due to the high coupling between the three equations, which makes the analytical study of these equations difficult, we solved this system numerically following a finite differences scheme. We impose homogeneous initial conditions for the three variables and run the simulation with a stop criterion satisfying stationary profiles to the three variables. Specifically, we use as stop criterion that the Norm-2 of difference between the profile of each variable and the previous one in the finite differences scheme is less than 0.01. The model was implemented using Matlab software.

**Figure 2.**
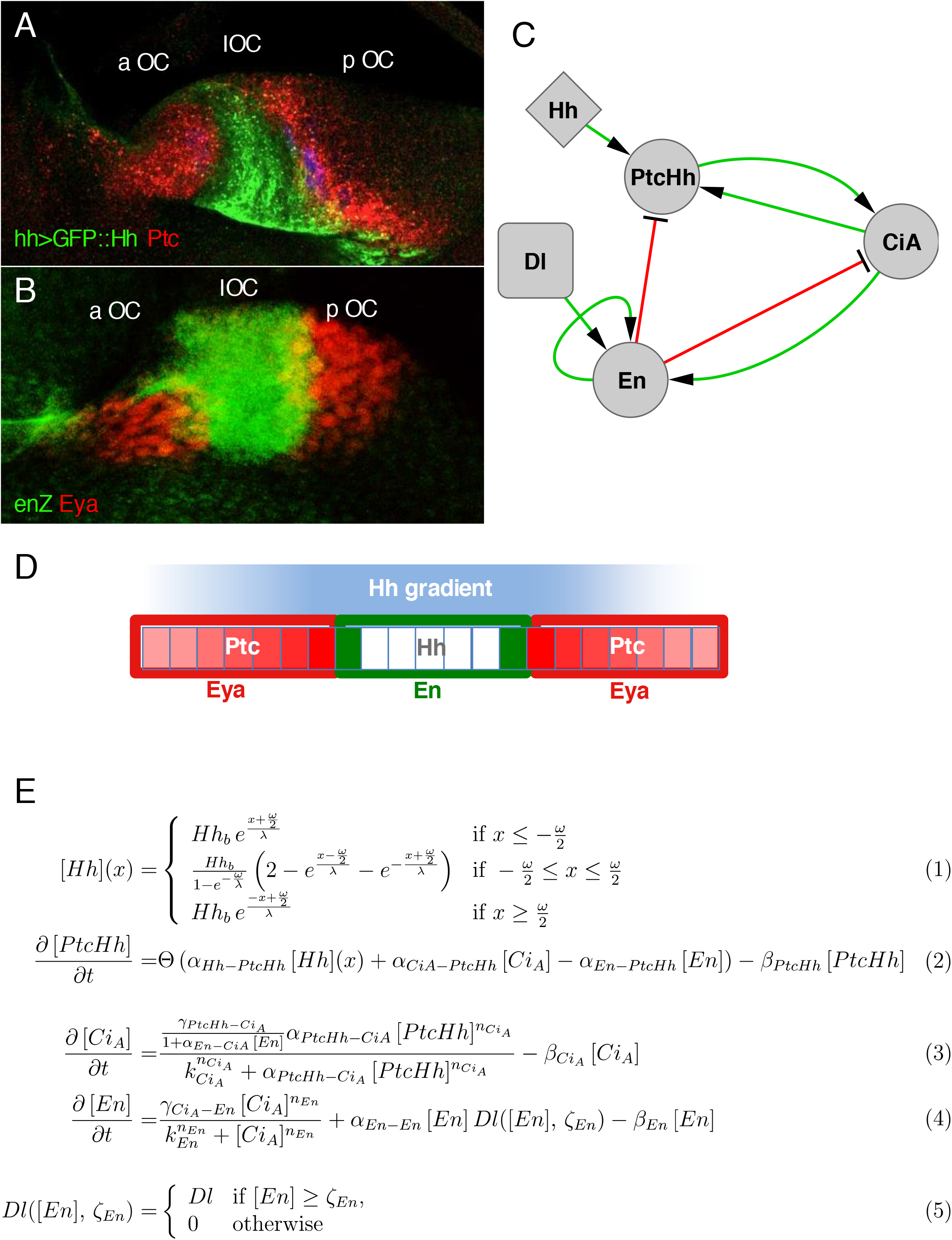
Patterning of the ocellar region primordium and the 3node-GRN. (A) Confocal image of an ocellar region primordium from a *hh-GAL4*, UAS-Hh::GFP *D. melanogaster* larva, stained for anti-GFP (Hh, green) and anti-Ptc (red). In this genotype Hh protein (Hh::GFP) is produced from its normal domain of expression (see Materials and Methods). (B) A similar ocellar region primordium from a *en-Z* larvae, stained for anti-β-galactosidase *(en-Z*, green) and Eya (red). aOC, pOC and IOC mark the anterior ocellus, posterior ocellus and interocellar prospective regions, respectively. (C) The 3node-GRN. Links are marked in green (activating) or red (repressive). (D) Schematic representation of the larval ocellar region as a monodimensional row of cells. Hh-producing cells (white) generate a time-invariant Hh gradient (blue). Ptc acts as Hh-signaling readout. Although initially widespread, the final pattern of Ptc (red) is in two domains adjacent to those of En (green). In the Ptc-domains, the Hh pathway remains active (i.e. there is expression of CiA, red) and the expression of Eya, a retinal determination gene, is established (not shown). The two outputs of the model are the lengths of the OC (marked by CiA) and of the IOC (marked by En) regions. (E) List of equations formalizing the 3node-GRN. Hh concentration at the boundaries of the source of width ω, centered at position x=0, reads Hh_b_ =α_Hh_/2β_Hh_(1-exp(ω/λ)). The model contains different parameter types: α_x_ for the basal transcription rates, β_x_ for the degradation rates, k_x_ for the Hill equation transcriptional regulators and n_x_ for the Hill coefficients. Subscript X-Y, with X and Y system variables, indicates a regulation from X to Y. For example, α_En-PtcHh_ is transcription rate parameter of the interaction from En to PtcHh. Parameter ζ_En_ indicates the En concentration threshold upon which En is self-regulated (see equations 4 and 5).

### Parameters sensitivity analysis and phenotypic phase space

To perform the parameters sensitivity analysis we run simulations in the model fixing all the parameters to a control value but one, which is randomized over two orders of magnitude around its control value. This process is repeated for each parameter. The resulting CiA pattern of the simulations (A) is compared to the pattern obtained by the control set of parameters (B). We measure the Euclidean distance (λ) between the two normalized patterns to obtain a *goodness* value for the randomized simulation (eq. 6).

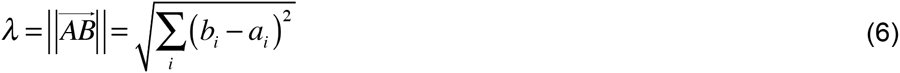

where *a_i_* and *b_i_* are the components of vectors A and B, respectively.

The distance distributions are shown in Fig. S1 (considered as complementary distance, 1-λ) for all the parameters. From this analysis we can extract important information about which parameters are more sensitive or more insensitive to variations away from the control parameter values. A complementary distance value of 0.8 was selected as a “goodness” threshold, as every pattern checked for a parameter set with a complementary distance value equal or higher to this value fits the target ocellar pattern. Following this “goodness” threshold, every parameter whose distance distribution falls below 0.8 is considered “sensitive”. We find that all the parameters in the simplified model can be considered as sensitive.

To evaluate whether the simplified model shows a restricted phenotype space of the ocellar region, we performed simulations (N=9000) with randomized parameters, modifying random seeds, within three goodness intervals 1-λ≥0.8 (‘good’), 0.8>1-λ≥0.6 (‘medium’) and 0.6>1-λ≥0.4 (‘bad’), (3000 simulations each) (Fig. 3A). Effective Hh diffusion coefficient D was varied in the following ranges: good=[0.068,0.109], medium=[0.068,0.010] and bad=[0.068,0.010] in μm^2^s^-1^. The effective turnover of Hh, β_Hh_, was varied in the following ranges: good=[2.1,2.5], medium=[1.5,2.1] and bad=[1.0,1.5] in 10^-4^s^-1^. Figure S2 shows two morphospace samples of the 3node-GRN (A) and including parameters D and β_Hh_ (B). Both samples contain 9000 points each with the same random seed.

**Figure 3.**
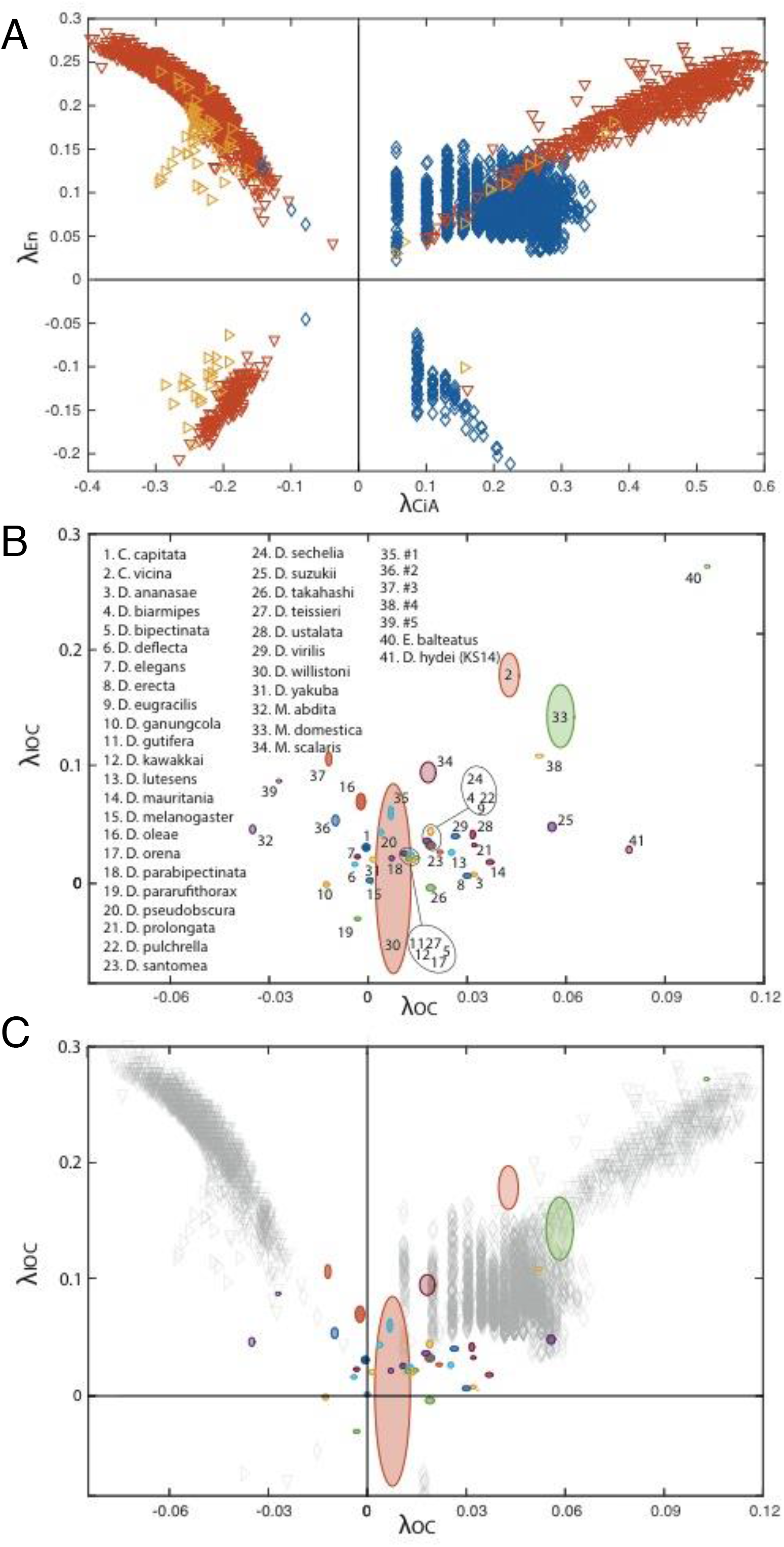
Predicted and measured phenotypic space. (A) Distribution of the phenotypic space defined by variations in simulated En profiles against variations in the simulated CiA profiles, to a profile marked as control. Different colors identify different parameter ranges for complementary distance in the parameter sensitivity analysis (see SFig 1 for a full parameter sensitivity analysis): 1-λ=0.8 (‘good’, blue), 1-λ=0.6 (‘medium’, red) and 1-λ=0.4 (‘bad’, yelow). (B) Distribution of IOC against OC sizes for a screening of 41 fly species. The ellipses defining each fly in the phenotypic space are centered in the average size value and their axes show the standard deviation from the mean. The colors of the ellipses show proportional opacity to the number of samples, varying from N=2 to N=14. The following species were not fully classified: #1 *Heleomyzidae* sp., #2 *Drosophila* sp., #3 *Apotropina* sp. #4 Schyzophora, Brachycera, Acalyptratae, with complete subcosta, costal break, arista almost bare, #5 Calyptratae, Muscoidea, Anthomyiidae. (C) Overlap between the phenotypic spaces for simulated and measured fly species.

### Phenotypic Classification using Bayesian Networks (BNs)

In this work, the same dataset of parameters is used to attempt the prediction of three types of phenotype class: OC size (OC), IOC size (IOC) and Near/Far (NF). Thus, three different classification problems are attempted with the same machine learning method. For each parameter set (instance) we calculated λ_CiA_, λ_En_ and (λ_CiA_^2^ + λ_En_^2^)^1/2^ for the class OC, IOC and NF respectively. In OC, values with λ_CiA_ <0 (λ_CiA_ > 0) received class value 0 (1). In IOC, values with λ_En_<0.15 (λ_En_ ≥ 0.15) received class value 0 (1). And in NF, values with (λ_CiA_^2^ + λ_En_^2^)^1/2^ < 0.3 ((λ_CiA_^2^ + λ_En_^2^)^1/2^ ≥ 0.3) received class value 0 (1). Learning takes place in the following way:

1. The instances in the dataset are divided in ten subsets. Each subset must have a collection of instances that is representative of the whole dataset.
2. Subset 1 is chosen as a Test subset, while the remaining subsets are used to train the learning method.
3. The machine learning method takes the training subset and infers the relationship between parameters needed to determine the phenotype class for every instance.
4. This learning process is then validated using the test subset, comparing the actual phenotype classes with the ones predicted by the machine learning method. The success rate (percentage of classes correctly predicted) is called predictive accuracy.
5. Steps 2-4 are repeated using each one of the 10 subsets as test subsets, while the other 9 subsets are used as training subsets in each case. This system of swapping subsets as tests is called 10-fold cross validation. It is used to increase the chances of having a representative test sample.
6. The 10 test predictive accuracies obtained from this repetition are averaged, giving a final predictive accuracy for this method, using this dataset.

A BN learns from the data provided by arranging the parameters in an ascending network, where the relative probability between parameters is established. Thus, the heuristics of BN returns a network of relative probabilities between parameters. Parameters are related to one another by probability distributions, according to the frequency (combined or not), with which a certain parameter has a certain value. For example, in order to establish the statistical relationship between a parent parameter and a child parameter, the question being asked is: provided that this child parameter (for this particular instance) has a certain value *X*, what range of values are expected on this parent parameter, and what probabilities are assigned to those ranges. These probabilities are expressed with the basic formula of Bayes’ Theorem:

Knowing:

- The frequency P(Y) with which a parent parameter has a value *Y.*
- The frequency P(X) with which a child parameter has a value *X*.
- The relative frequency P(X|Y) with which, having *Y* in the parent parameter, we have *X* in the child parameter.

We can obtain the relative frequency P(Y|X) with which, having *X* in the child parameter, we have *Y* in the parent parameter, according to:

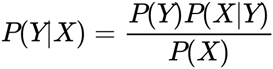

As we climb up the network of parameters, these ranges and probabilities are refined in accordance to an optimal classification, thus maximizing the predictive accuracy. The final inference is made from the topmost parameter or parameters, to the class. This is when the network class is decided. The connections in these networks are averaged in one final network that represents the overall connections of the parameters of a certain dataset, needed to correctly classify instances. Since, in this work, the aim is to classify three different phenotypic classes using the same set of parameters, it follows that three networks (one per classification problem) were obtained from the BN method.

A BN, as represented in this work (see Figure 4B), is read from the bottom up. At the top of the network lies the phenotype class. Parameters on higher levels of the network are considered as parents of the parameters immediately below. Parameters are related to one another by probability distributions, according to the frequency (combined or not), with which a certain parameter has a certain value (or lies within a certain interval). As we climb up the network of parameters, the intervals and probabilities are refined in accordance to an optimal classification, thus maximizing the predictive accuracy. The final inference is made from the topmost parameter or parameters, to the class. This is when the class is decided. Classification with Bayesian Networks was performed using WEKA 3.7.11 (Hall, 2009).

**Figure 4.**
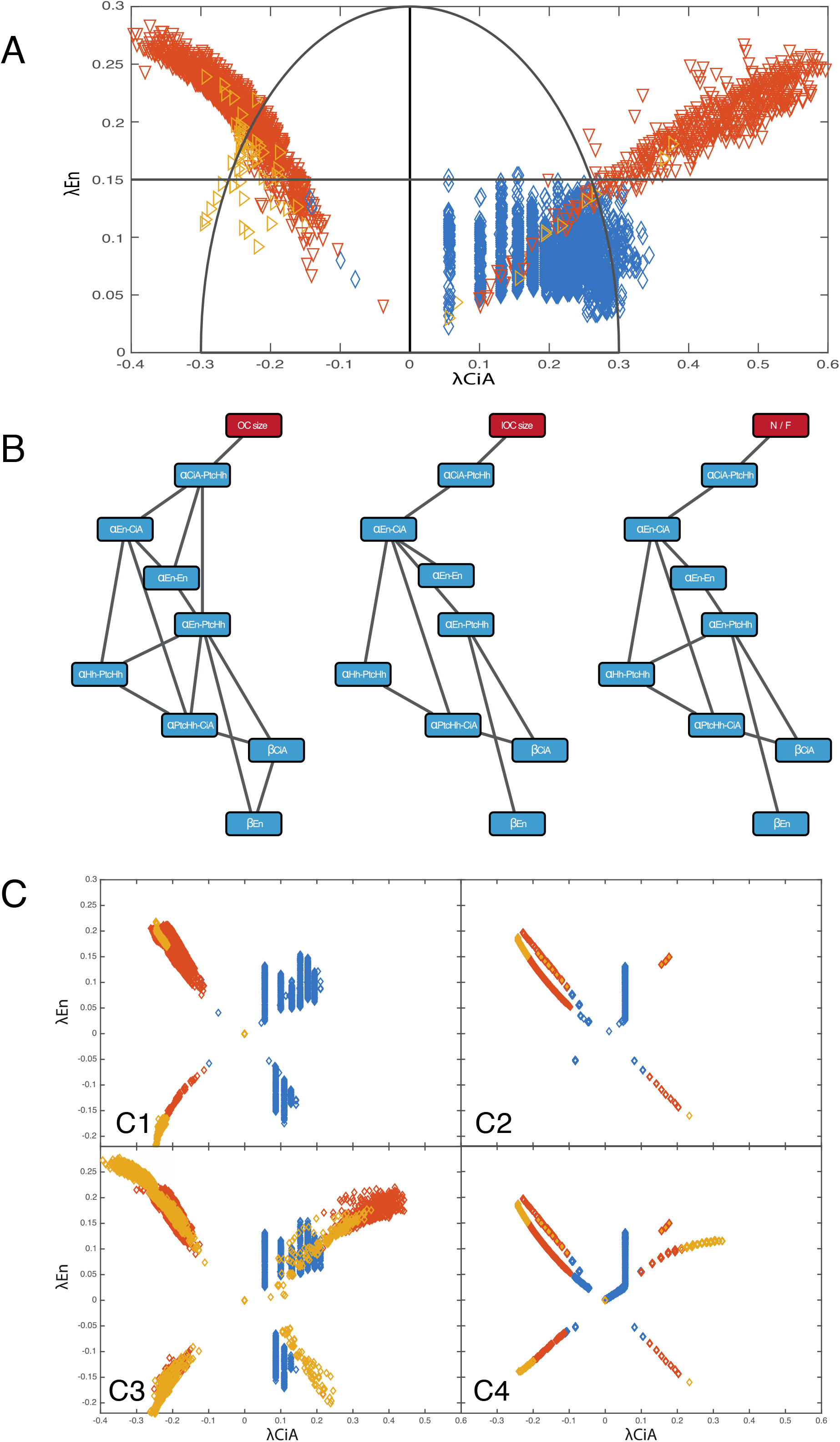
Bayesian Network (BN) analysis. (A) Classification of the phenotypic subspace (with λ_En_>0) according to the distance of the OC (λ_CiA_) and IOC (λ_En_) lengths to the control. Horizontal line at λ_En_=0.15 delimits zones in the IOC size morphological subdivision. The phenotypes within the semicircle are considered similar (“near”) to the control value, while those outside the semi-circumference (centered at (λ_CiA_=0, λ_En_=0) and radius of 0.3) are considered less so (“far”). (B) Bayesian networks for the three phenotypic classes: large/small OC (left), large/small IOC (middle) and ocellar regiones near/far from the control N/F (right). These networks show eight parameters with the greatest influence to define phenotypic space localizations. (C) Simulations of the 3node-GRN randomizing the three top-most parameters predicted by the BN simultaneously (C1) and independently (C2), and including parameter β_En_ in a simultaneous (C3) and independent (C4) randomization.

## RESULTS

### A simplified GRN model recapitulates the ocellar pattern and predicts a specific morphological space for the ocellar region

The ocellar region (Figure 1A) arises from the fusion along the dorsal midline of the left and right cephalic primordia (often called “eye-antennal imaginal discs”; Figure 1B). In each primordium, a single Hedgehog (Hh)-producing domain provides cells with positional information, by generating a signaling gradient. Signaling activity can be visualized using the expression levels of *patched* (Ptc) as its readout (Figure 2A,D). This is so because Ptc, in addition to being the Hh receptor, is a positive target of the pathway –i.e. the levels of Ptc increase as the signal intensity increases (Chen and Struhl, 1996). Activation of the Hh pathway leads to the stabilization of the activator form of *cubitus interruptus* (CiA), the Gli-type transcription factor that mediates the nuclear transduction of the pathway (Alexandre et al., 1996). The Hh signaling gradient is then translated into two cell fates. At its highest levels, and basically coinciding with the Hh-producing cells, the pathway activates the expression of the transcriptional repressor *engrailed* (En). This leads to a pathway shut off, as En represses the transcription of *ci* and *ptc.* This signaling-Off region gives rise to the interocellar cuticle (IOC). Maintenance of En expression in the IOC region requires *Delta (Dl)/Notch* signaling (Aguilar-Hidalgo et al., 2013). Flanking the Hh-producing/En-expressing domain, graded Hh signaling results in the stabilization of CiA which, in turn, activates the expression of genes that specify the ocellar retinas, including *eyes absent (eya)* (Blanco et al., 2009) on both sides of the Hh-producing domain (Figure 2B). During metamorphosis, as the two cephalic primordia fuse, the two anterior Eya domains merge into the anterior (or *medial*), unpaired, ocellus (aOC), while the two posterior domains remain separate and form the two posterior (or *lateral)* ocelli (pOC). As mentioned above, the region in between the two Eya patches expresses the transcription factor *engrailed* (*en*) and form the intervening interocellar cuticle (IOC) in the adult (see Figure 1B). Therefore, the early patterning of the ocellar region entails the generation of basically two cell fates (OC and IOC), the control of their respective size, and their spacing into an “OC-IOC-OC” pattern.

As mentioned, the evolutionary conserved Hedgehog (Hh) signaling pathway plays a pivotal role in these processes of fate assignment and size control. Although the pattern is bidimensional, it can be simplified as a monodimensional process along the anteroposterior axis (Aguilar-Hidalgo et al., 2013), and described by two variables, the lengths of the OC and the IOC distance (schematized in Figure 2D). A previous model of the detailed GRN, including 13 molecules (such as CiA) or molecular complexes (such as Ptc:Hh) as network’s nodes, predicted that the phenotypic space available to the GRN (i.e. the sets of OC and IOC lengths) was limited. This being so, the analysis of the model could identify the parameter, or subset of parameters with the largest impact on size variation. However the size and complexity of the model makes this analysis difficult. To make this analysis more tractable, we resorted to a simplified GRN model that retains critical genetic/molecular interactions and which we showed previously that recapitulates the ocellar pattern (Aguilar Hidalgo et al., 2015)(see Figure 2C). Pattern in this GRN is dependent on the specific topology of a core regulatory network motif containing an activator-repressor regulatory mechanism describing the dynamics of 3 variables with 16 parameters, what we call the “3-node GRN” (Aguilar Hidalgo et al., 2015). We solved the model to find the steady state pattern –i.e. that is the final, stable pattern that is reflected in the adult ocellar complex. As the equations of the “3-node GRN” contain nonlinear terms, we chose to solve these numerically. Hh (equation 1) then serves as source for PtcHh complex production (Ptc being Hh receptor, equation 2), which activates the production of CiA (equation 3). CiA favors the maintenance of PtcHh and can activate expression of En (equation 4) and Eya, the two readouts of the model. En is a low-sensitivity Hh target and a repressor of the pathway components CiA and PtcHh (and therefore, of Eya). Above a certain concentration threshold ζ_En_, En is self-maintained (genetically, this step requires the Dl/Notch pathway (Aguilar-Hidalgo et al., 2013), equations 4 and 5) and becomes independent on the Hh signaling. Due to En being a low sensitivity target, En is only self-maintained in the zone of maximal Hh concentration that closely corresponds to the Hh-producing domain. The En-expressing domain gives rise to the IOC region. In regions adjacent to the Hh-producing domain, where the Hh concentrations are not enough as to activate En, CiA is stabilized and Eya expression is induced, generating the OC domains. Because Eya expression is induced by CiA, in the model CiA is used as a marker of OC identity. Therefore, the variables that define the morphology of the ocellar complex are lengths of the En and CiA domains, which represent the IOC and OC regions, respectively.

In order to find the parameters for which small variations caused significant deviations from “control” OC and IOC lengths (see methods), that represents *D. melanogaster*, we first performed an individual sensitivity analysis for each of the 16 3node-GRN parameters. To find a metric for this deviation we calculated the distance between the control pattern and the patterns generated by varying each of the parameters. We established three thresholds for the complementary of this distance (1-λ): 1-λ≥0.8 (‘good’), 0.8>1-λ≥0.6 (‘medium’) and 0.6>1-λ≥0.4 (‘bad’), with 1-λ≥0.8 giving the patterns closest to the control. This analysis showed that every parameter in the 3-node GRN is sensitive to small variations, as their distributions mostly fall below the 0.8-threshold (Figure S1). Then, we performed simulations using randomized values (from the ‘good’, ‘medium’ and ‘bad’ intervals) for every parameter simultaneously to generate a point (a “phenotype”) in the phase space. Therefore, this phase-space is a ‘phenotype space’ or ‘morphospace’. The axes of this phenotype space represent the deviations of the lengths of the CiA and En expression domains (λ_CiA_, λ_En_) from the control (at (0,0)). For example, (-0.20, 0.25) would be an ocellar complex with smaller OC (λ_CiA_= -0.20) and larger IOC (λ_En_= +0.25) than the control. We found that: (1) The simulations with randomized parameter sets show a non-random distribution, yielding a sort of “butterfly wing” pattern in the phenotype space (Figure 3A); In addition, (2) the model may yield very similar phenotypes even when the randomized parameters come from different goodness intervals (i.e. the results, expressed as a point (λ_CiA_, λ_En_) in the morphospace, lie close to one another) (Figure 3A). (3) However, we also find that the “goodness” of parameters biases the distribution of solutions in the morphospace. Thus, parameter values chosen from the “good” interval mostly result in larger OC than the control (i.e. positive λ_CiA_), while “medium” and “bad” parameter values avoid larger OC AND smaller IOC values. In addition, globally considered, parameter variation in the 3node-GRN tends to yield ocellar regions with larger IOC (i.e λ_En_>0) (Figure 3A). Although our study focuses on the intracellular GRN driving the ocellar pattern, we analyzed to what extent the variation of parameters affecting the gradient of Hh affected the shape of the morphospace. Specifically, we varied the effective Hh diffusion coefficient D and the effective turnover of Hh, β_Hh_, as these parameters together define the gradient’s length scale λ=(D/β_Hh_)^1/2^ (see Eq. 1). We found that the extended morphospace that resulted distributed “medium” and “bad” parameter spreads slightly further away from control values. However, globally, the extended morphospace is very similar to the 3-node GRN’s with a fixed Hh gradient (Figure S2). Therefore, the intracellular GRN determines, to a great extent, the ocellar complex phenotype space. In what follows we continue our analysis of the intracellular 3-node GRN without considering variations in the extracellular Hh gradient.

### Quantitative phenotypic variation of the ocellar region in different fly species

The study of the phenotype space allowed by the 3-node GRN predicted that simultaneous variation of all parameters (by assigning each parameter a random value within a certain interval; see Methods) should result in non-random phenotypes –i.e. the phenotypic space available for morphological variation is limited. To test whether this prediction agrees with the phenotypic variation observed in actual fly species, we measured the length of the anterior and posterior OC and the IOC distance in a sample of 41 fly species (Figure 3B). To account for body size differences, these measurements were normalized using the inter-anterior occipital bristle distance, as a proxy of head width. Only females were measured. The species set surveyed is not comprehensive across Schizophoran flies and is strongly biased towards Drosophilidae species close to *Drosophila melanogaster*, for which we had the easiest access to (see Materials and Methods). When plotted, the distribution of (λ_OC_, λ_IOC_), which represents the variation in the respective OC and IOC lengths (only pOC were used) relative to *D. melanogaster*, showed a pattern resembling the “butterfly wing” pattern predicted by the model (Figure 3B).

In general, we find that species belonging to groups far away from Drosophilidae show the most divergent morphologies. Such is the case of *Megaselia abdita* (Phoridae, no. 32 in Figure 3B), *Episyrphus balteatus* (Syrphidae, no. 40 in Figure 3B), or *Musca domestica* (Muscidae, no. 33 in Figure 3B). This qualitative similarity in distributions is best observed when the predicted and measured phenotypic spaces are overlapped (Figure 3C). Although the similarity noticed is purely qualitative and based on a limited sample of species, and therefore still has to be regarded as preliminary, we find it lends support to the idea that, in nature, the phenotypic variability available to the ocellar region is also restricted and follows similar patterns as those predicted by the model.

### Machine learning method finds parameter relations defining ocellar and interocellar sizes

For each parameter set, the 3-node GRN yields a value for the OC and IOC lengths –i.e. defines a point in the ‘phenotypic space’. But does every parameter contribute equally to localize a point in this space or, instead, one parameter (or a subset of parameters) has a major contribution to determining the localization of this point -that is to morphological *variation?* If the latter were the case, the identification of this set of *control parameters* may point to genetic/molecular links of particular relevance in controlling the OC and IOC lengths.

In order to establish a relationship between the parameters in the 3-node GRN and the morphological variation of the ocellar region we can envision a number of potential approaches. A developmental genetics approach, without prior knowledge, would entail the systematic perturbation of the genetic links implicit in the 16 parameters of the model, alone and in combination. A quantitative genetics approach (QTL) would be capable of identifying important elements of the network, but it would be limited to cross-hybridizing species showing significant differences in ocellar morphology. In addition, a QTL approach could be capable of identifying causes for existent variation, not for all potential variation. From a numerical perspective, the full parameter space is vast. An alternative to dynamical model simulation analysis could be the use of classification methods to infer morphological variation directly from the randomized parameter vectors. One such method is Bayesian Networks (BNs) (Pazzani, 1996; Friedman et al., 1998; Keogh and Pazzani, 1999). A Bayesian Network (BN) is an acyclic, directed graph connecting a series of variables linked by their conditional probabilities (non-linked variables are independent from each other). These BNs can be used to compute the probability of a given output. In our case, the variables are the 13 parameters of the 3-node GRN, and the output is whether a “phenotype” (a point in the (λ_CiA_, λ_En_) plane) falls within a given region of this space. As we climb up the network of parameters, the conditional probabilities maximize the predictive accuracy (for a more detailed description of the BN learning method and classification, please see Methods). Specifically, we used this method to try to identify relevant parameters for morphological variation.

We subdivided the phenotypic space into three different morphological classes: (1) OC smaller or larger than the control (Left: λ_CiA_<0 or Right: λ_CiA_>0, respectively); (2) small or large IOC (Up: λ_En_<0.15 or Down: λ_En_≥0.15, respectively). And (3) “Near/Far” (N/F), which distinguishes between positions in the phenotypic space that are more or less similar (“near” or “far”, respectively) to the control. In this case we impose the same sign to the size variation of the OC and IOC -that is, large OC with large IOC, and small OC with small IOC. Specifically, a point is “near” the control (i.e. it is “similar”) if it is located inside a circumference with radius 0.3. If the point is located outside the circumference, it is classified as “far” from the control (see Figure 4A). Note that we consider only points with λ_En_≥0 due to the low number of points with λ_En_<0 (i.e. the model does not yield many cases of ocelli smaller than the “control”). We applied BN analysis to identify parameters which, when co-varied, localize points to one of these zones. For each class, the BN heuristics returned a network of relative probabilities between parameters, with very good classification results (90.35% for N/F, 96.23% for OC, and 94.58% for IOC). The analysis of the three networks, that establish a hierarchy of relations between parameters (in Figure 4B the networks includes the set of 8 parameters with the highest classification value), resulted in a number of observations. First, the three BNs show the same nodes in a similar hierarchy, despite the fact that they inform about different phenotype classes. This implies that the same genetic interactions (represented by parameters in the model) control the variation of different phenotypic classes. Second, the three top-most parameters in each BN suffice for a good classification. These three parameters include, with decreasing relevance, the one determining the transcriptional efficiency by which CiA activates Ptc expression (α_CiA-PtcHh_), the intensity of repression of CiA by En (α_En-CiA_), and α_En-En_, that controls *en* autoregulation.

To validate the BN results, we compared the morphospace generated when the three predicted control parameters (α_CiA-PtcHh_, α_En-CiA_ and α_En-En_) were randomly co-varied with the morphospace resulting from the overlap of the three simulations generated when each of the parameters were varied individually. While the morphospace resulting from parameter co-variation recapitulated most of the butterfly wing pattern (Figure 4C1), the ones resulting from varying the parameters individually matched the butterfly wing pattern much more poorly (Figure 4C2). Still, covariation of the three top-ranked parameters missed the “right forewing” (i.e. λ_CiA_>0, λ_En_>0). We sought among the five remaining parameters in the BNs the parameter or parameters, that when co-varied, showed the missing “wing”. We found that β_En_, which correspond to the degradation rate of En, when co-varied with the three top parameters in the BNs yielded the “butterfly wing” pattern (Figure 4C3). Again, this pattern was just sketched when the four parameters were independently randomized and their patterns overlapped (Figure 4C4). This analysis indicates that the control of morphological variations in the ocellar region requires the cooperation of four major parameters. In addition, we noted that, of the 16 parameters, those corresponding to non-linear terms in the model, such as Hill coefficients, have the least relevance in the classification in the three BNs (OC, IOC and N/F) (not included in the BNs in Figure 4). Finally, although similar, the exact topology of the three networks varies, with the BN for OC size being the most connected.

## DISCUSSION

In this paper we have studied the ocellar GRN, as an example of gene network regulated by the Hh morphogen, to predict the range of available phenotypic space for morphological variation and tried to predict parameters within this network with a major effect in controlling that morphological variation. We have found that a simple 3-node GRN that recapitulates the pattern of the ocellar region predicts restrictions to variations in the size of the ocelli (OC) and the distance in between the ocelli (IOC). When measured, the distribution of OC and IOC lengths from a sample of dipteran species seemed to follow, qualitatively, the same distribution in the phenotypic space that the one predicted by the model. We take this result as lending support to the notion that the GRN structure indeed restricts the evolvability not only of the model’s output, but also of its real surrogate, as these restrictions would be reflected by the actual phenotypes found in nature. However, as we noted, this conclusion is tentative. First, because the sample of species is not sufficiently large and comprehensive across the higher dipterans. Second, because the morphologies in extant species may as well be the result of natural selection –i.e. the pattern of morphologies observed having been shaped by functional constraints, such as ocellar regions having an IOC length above a certain limit, to allow the aOC and pOC to scan separate regions of vision (however, for this particular example, we note that the model also predicts that too short IOC distances are unlikely). We believe that most likely the actual phenotypes have resulted from the action of natural selection of the advantageous phenotypes from the morphospace allowed by the GRN’s structure.

To more precisely define the contribution of gene regulatory steps to shaping the ocellar morphospace we envision two approaches. A developmental genetics approach in which, by using a priori information of the most likely relevant parameters, the morphological variation of allelic series in genes affecting those parameters is used to compare the predicted to the actual phenotypes measured in each allelic combination. A second approach would be a comparative one: to increase the size and breadth of the sample of dipteran species studied to examine how closely their ocellar morphologies map within the predicted “butterfly”-shaped morphospace, so that the closer the correlation, the more likely that the phenotypic range is determined by the GRN structure.

Basic to our approach to studying the role that gene network structure has in controlling the evolvability of the ocellar region (as a model of a Hh-patterned organ) is the assumption that *the GRN structure remains constant* in the species we examine. This allows us to compare different morphologies generated by *the same GRN* structure through the sole quantitative variation of its parameters. Although this assumption may seem a strong one, we think it justified. The 3-node GRN comprises a set of Hh-related regulatory linkages that have been shown to be operating in other developmental contexts, including a Hh source and a steady state Hh gradient; the basic Hh signal transduction path hh→Ptc:Hh→CiA→Ptc:Hh; or the CiA→En-ICiA repression feedback. This likely also extends to the activation of retinal genes, such as *eya*, by the Hh signaling pathway –i.e. they can be considered conserved regulatory modules, or “kernels” (Davidson et al., 2003), and therefore they are likely to be invariant in the network. Even, if new nodes were to appear during evolution, it is conceivable that their effect could be incorporated as a quantitative variation of some of the parameters that define the network. For example, recent work (Dominguez-Cejudo and Casares, 2015) has shown that the Six3-type transcription factor *Optix* is expressed in the aOC, and not in the pOC during development in *D. melanogaster.* During larval development the aOC primordium is smaller than the pOC primordium (DGM, FC, unpublished). One hypothesis is that Optix would modify some OC-controlling parameters in the network leading to a smaller sized-aOC. If this were the case, *Optix’s* action could be modeled implicitly as the variation of one parameter (specifically affecting the aOC) without the need to add it explicitly to the network model. Therefore, the network would still be of use to explore the potential range of morphologies even if not containing *explicitly* all the playing genes and interactions, provided that these elements and interactions can be represented implicitly in the model equations, and that they do not alter the 3-node network’s structure. (Note that our 3-node model is symmetrical –i.e. does not consider potential regulatory differences between anterior- and posterior OC). We have circumscribed our analysis to dipterans as we can more confidently assume the conservation of the GRN structure. Whether this model is applicable to other insects depends on whether the ocellar GRN is conserved beyond dipterans in these groups.

In principle, one of the advantages of the use of models is the possibility to extract information relevant to the behavior of the biological process modeled. If we accept the assumption that the GRN structure remains constant within higher Diptera (see above), an important point is to determine how parameter variation impacts morphological variation. The parameters in the model are surrogates of biochemical rate constants, including those for protein-protein interactions (i.e. activation of the Ptc receptor (as PtcHh) by its ligands), protein degradation and, most importantly, activating or repressing protein-DNA interactions between transcription factors and cis-regulatory elements. As sequence variation is generated in a given population, a mixture of variants will be combined in each individual of this population. Therefore, it is of interest to analyze the combined effects of allelic variants (i.e. parameter variants), rather than of individual variants, on the final morphology of the system. Even in our relatively simple 3node-GRN, a comprehensive analysis of parameter co-variation entails long calculations. Although doable, we have opted to introduce an alternative approach: the use of Bayesian Networks to identify the most relevant parameters in defining a particular morphological class and their probabilistic relationship. This approach has been recently used to identify critical interleukins within the murine cytokine-hormonal network (Field et al., 2015). In our BN analysis, four parameters stand out as most relevant: α_CiA-PtcHh_, α_En-CiA_, α_En-En_ and β_En_. The first three are transcriptional regulatory steps. α_CiA-PtcHh_ represents the activation rate of Ptc (which engages with Hh in an active PtcHh signaling complex) by the activator form of the Gli transcription factor *ci*: CiA. α_En-CiA_ reflects the repressing action of En on *ci* transcription (represented in the model as CiA repression), a regulatory step that controls the establishment of the IOC; and α_En-En_, which maintains the IOC region in the CiA-repressed region. We propose that these parameters, jointly, may be responsible for most of the morphological variation seen in the ocellar region in different species.

Another observation derived from the BN analysis is that variation in OC length is defined, at least probabilistically, by a more connected network than for the IOC length. This suggests to us that morphological variation of OC size is genetically more complex than that of the IOC. The “N/F” BN shows an intermediate complexity, as it reflects the phenotypic co-variation of OC and IOC. Finally, we noted that the eight parameters with significant contribution to defining morphological classes were linear terms in our model. The non-linear terms, that include, for example, the Hill constants have been shown to be required for the system’s stability (Aguilar Hidalgo et al., 2015). Therefore, from a modeling perspective, morphological variation is basically defined by the linear terms (transcriptional activations and repression and decay constants).

This study, combining GRN modeling and machine learning with biological measurements, indicates that morphological variation in the ocellar region is limited by the specific topology of its GRN and identifies a very short list of biochemical parameters, mostly representing transcriptional regulatory steps, that jointly control such variation. These results reinforce the notion that, as a general principle, the potential for morphological variation of organs is limited by the specific regulatory interactions governing their development, and that morphological variation can be the results of combination of genetic variants that modify, simultaneously, several biochemical parameters within those interactions.

## ACKNOWLEDGEMENTS

We thank J. Culi, B. Prud’Homme, P. Simpson, J. Jaeger, K Wotton and M. Averof for flies, and D. G. Míguez and M. Popovic for critical reading of the manuscript. Research at the Casares laboratory is funded by the Spanish Ministry for Economy and Innovation (MINECO) and Feder Funds through grant BFU2012-34324 to FC. DGM is a MINECO PhD Fellow. DBA was supported in part by the Spanish Inter-Ministerial Commission of Science and Technology under Project TIN2014-54583-C2-1-R, the European Regional Development fund, and the “Junta de Andalucía” (Spain), under Project P2011-TIC-7508. We also thank A. Iannini for technical assistance, the Developmental Studies Hybridoma Bank, University of Iowa, for antibodies and the CABD Advanced Light Microscopy Facility for help with confocal imaging and members of the Casares lab for discussions.

---

**Figure S1.**
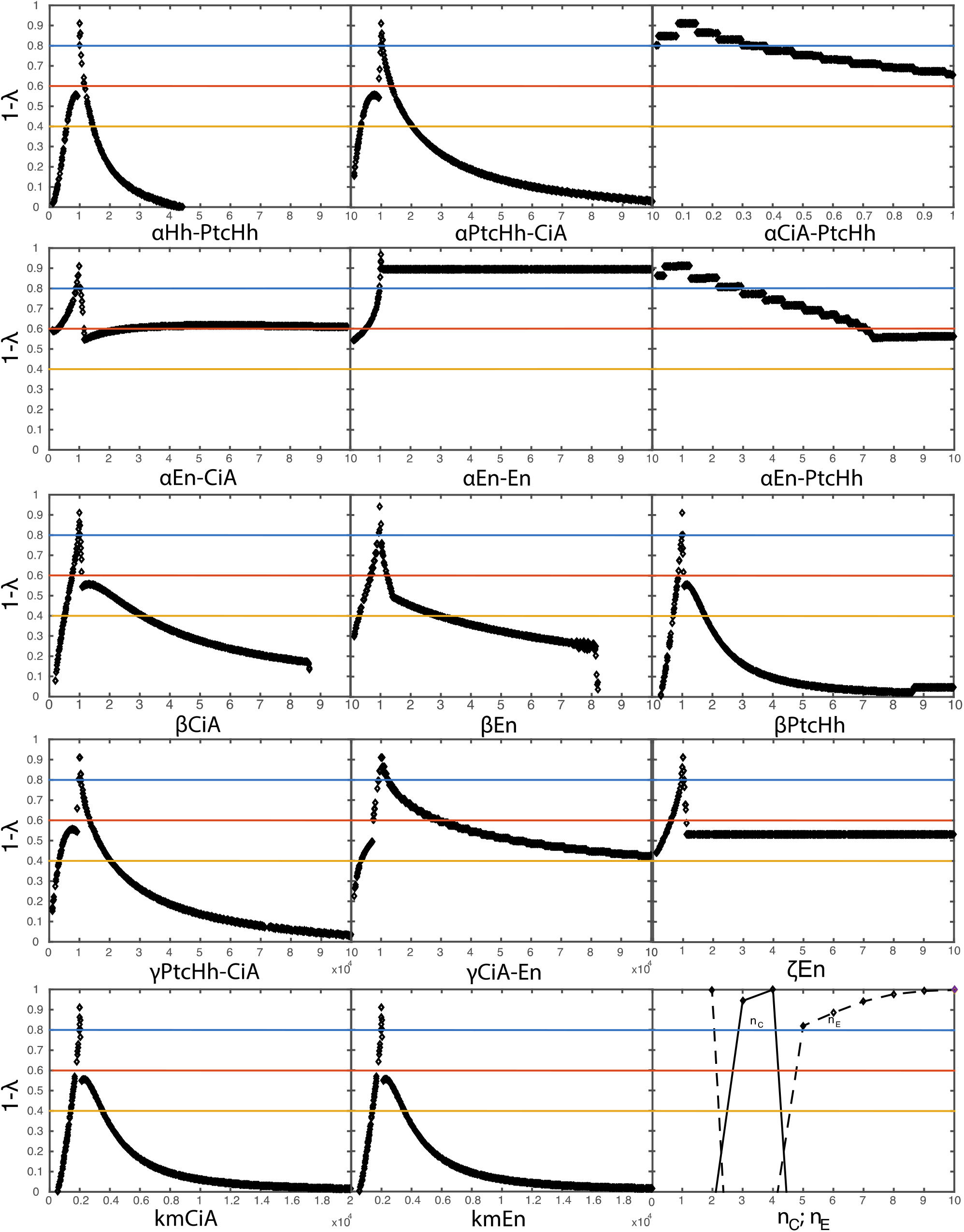
Aguilar-Hidalgo et al.

**Figure S2.**
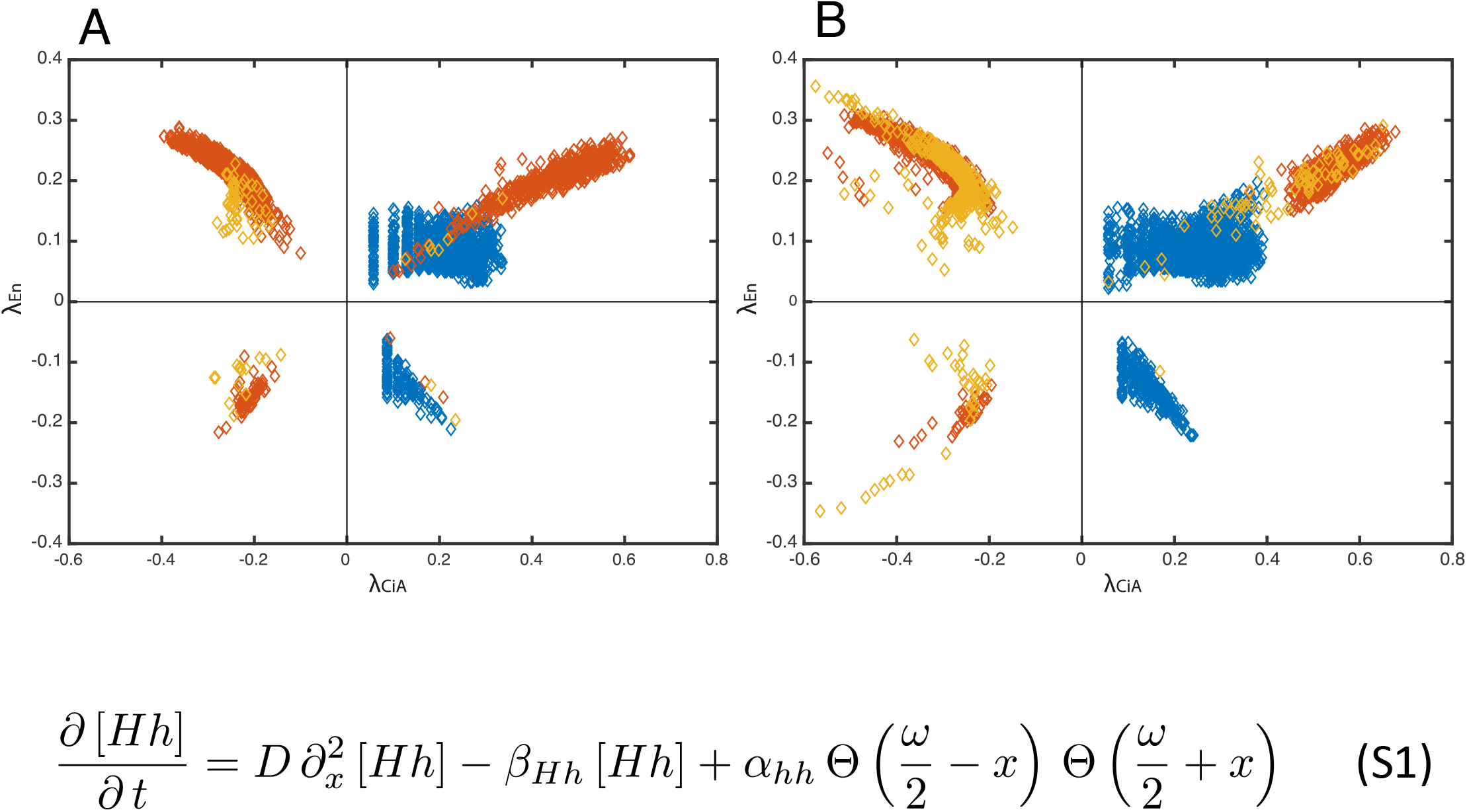
Aguilar-Hidalgo et al.

## REFERENCES

Aguilar Hidalgo, D., Lemos, M. C. and Cordoba, A. (2015) ‘Core regulatory network motif underlies the ocellar complex patterning in Drosophila melanogaster’, Physica D: Nonlinear Phenomena 295-296: 91–102.

Aguilar-Hidalgo, D., Dominguez-Cejudo, M. A., Amore, G., Brockmann, A., Lemos, M. C., Cordoba, A. and Casares, F. (2013) ‘A Hh-driven gene network controls specification, pattern and size of the Drosophila simple eyes’, Development 140(1): 82–92.

Alexandre, C., Jacinto, A. and Ingham, P. W. (1996) ‘Transcriptional activation of hedgehog target genes in Drosophila is mediated directly by the cubitus interruptus protein, a member of the GLI family of zinc finger DNA-binding proteins’, Genes Dev 10(16): 2003–13.

Arnone, M. I. and Davidson, E. H. (1997) ‘The hardwiring of development: organization and function of genomic regulatory systems’, Development 124(10): 1851–64.

Arthur, W. (2006) Biased Embryos and Evolution: Cambridge University Press.

Blanco, J., Seimiya, M., Pauli, T., Reichert, H. and Gehring, W. J. (2009) ‘Wingless and Hedgehog signaling pathways regulate orthodenticle and eyes absent during ocelli development in Drosophila’, Dev Biol 329(1): 104–15.

Brockmann, A., Dominguez-Cejudo, M. A., Amore, G. and Casares, F. (2011) ‘Regulation of ocellar specification and size by twin of eyeless and homothorax’, Dev Dyn 240(1): 75–85.

Callejo, A., Culi, J. and Guerrero, I. (2008) ‘Patched, the receptor of Hedgehog, is a lipoprotein receptor’, Proc Natl Acad Sci U S A 105(3): 912–7.

Casares, F. and Mann, R. S. (2000) ‘A dual role for homothorax in inhibiting wing blade development and specifying proximal wing identities in Drosophila’, Development 127(7): 1499–508.

Rasband, W.S., ImageJ, U. S. National Institutes of Health, Bethesda, Maryland, USA, http://imagei.nih.gov/ij/, 1997–2015.

Chen, Y. and Struhl, G. (1996) ‘Dual roles for patched in sequestering and transducing Hedgehog’, Cell 87(3): 553–63.

Davidson, E. H. and Erwin, D. H. (2006) ‘Gene regulatory networks and the evolution of animal body plans’, Science 311(5762): 796–800.

Davidson, E. H., McClay, D. R. and Hood, L. (2003) ‘Regulatory gene networks and the properties of the developmental process’, Proc Natl Acad Sci U S A 100(4): 1475–80.

Dominguez-Cejudo, M. A. and Casares, F. (2015) ‘Anteroposterior patterning of Drosophila ocelli requires an anti-repressor mechanism within the hh pathway mediated by the Six3 gene Optix’, Development 142(16): 2801–9.

Felix, M. A. (2012) ‘Evolution in developmental phenotype space’, Curr Opin Genet Dev 22(6): 593–9.

Field, S. L., DAsgupta, T., Cummings, M. R., Savage, R. S., Adebayo, J., McSara, H., Gunawardena, J. and Orsi, N. M. (2015) ‘Bayesian modeling suggests that IL-12 (p40), IL-13 and MCP-1 drive murine cytokine networks in vivo’, BMC Systems Biology 9: 76.

Friedman, N., Goldszmidt, M. and Lee, T. J. (1998) ‘Bayesian Network Classification with Continuous Attributes: Getting the Best of Both Discretization and Parametric Fitting.’, ICML 98: 179–187.

Hall, M. (2009) ‘The WEKA data mining software: an update’, ACM SIGKDD explorations newsletter 11(1): 10–18.

Harjunmaa, E., Seidel, K., Hakkinen, T., Renvoise, E., Corfe, I. J., Kallonen, A., Zhang, Z. Q., Evans, A. R., Mikkola, M. L., Salazar-Ciudad, I. et al. (2014) ‘Replaying evolutionary transitions from the dental fossil record’, Nature 512(7512): 44–8.

Jaeger, J. and Monk, N. (2014) ‘Bioattractors: dynamical systems theory and the evolution of regulatory processes’, J Physiol 592(Pt 11): 2267–81.

Jaeger, J., Surkova, S., Blagov, M., Janssens, H., Kosman, D., Kozlov, K. N., Manu, Myasnikova E., Vanario-Alonso, C. E., Samsonova, M. et al. (2004) ‘Dynamic control of positional information in the early Drosophila embryo’, Nature 430(6997): 368–71.

Kauffman, S. A. (1993) The origins of order. Oxford University Press.

Keogh, E. and Pazzani, M. (1999) Learning Augmented Bayesian Classifiers. A Comparison of Distribution-Based and Classification-Based Approaches Seventh International Workshop on Artificial Intelligence and Statistics.

Lopez-Rios, J., Duchesne, A., Speziale, D., Andrey, G., Peterson, K. A., Germann, P., Unal, E., Liu, J., Floriot, S., Barbey, S. et al. (2014) ‘Attenuated sensing of SHH by Ptch 1 underlies evolution of bovine limbs’, Nature 511(7507). 46–51.

Oster, G., Shubin, N., Murray, J. D. and Alberch, P. (1988) ‘Evolution and morphogenetic rules. the shape of the vertebrate limb in ontogeny and phylogeny’, Evolution 42(5). 862–884.

Pazzani, M. J. (1996) Searching for Dependencies in Bayesian Classifiers. Springer.

Raspopovic, J., Marcon, L., Russo, L. and Sharpe, J. (2014) ‘Modeling digits. Digit patterning is controlled by a Bmp-Sox9-Wnt Turing network modulated by morphogen gradients’, Science 345(6196). 566–70.

Royet, J. and Finkelstein, R. (1996) ‘hedgehog, wingless and orthodenticle specify adult head development in Drosophila’, Development 122(6). 1849–58.

Royet, J. and Finkelstein, R. (1997) ‘Establishing primordia in the Drosophila eyeantennal imaginal disc. the roles of decapentaplegic, wingless and hedgehog’, Development 124(23). 4793–800.

Salazar-Ciudad, I. and Jernvall, J. (2010) ‘A computational model of teeth and the developmental origins of morphological variation’, Nature 464(7288). 583–6.

Waddington, C. H. (1957) The Strategy of the Genes: A Discussion of Some Aspects of Theoretical Biology, London. Ruskin House/George Allen and Unwin Ltd.

Wiegmann, B. M., Trautwein, M. D., Winkler, I. S., Barr, N. B., Kim, J. W., Lambkin, C., Bertone, M. A., Cassel, B. K., Bayless, K. M., Heimberg, A. M. et al. (2011) ‘Episodic radiations in the fly tree of life’, Proc Natl Acad Sci U S A 108(14). 5690–5.

